# Differential gene expression and gene-set enrichment analysis in Caco-2 monolayers during a 30-day timeline with Dexamethasone exposure: A data modelling approach to understanding culture age as co-variate for differential expression in a non-renewing epithelial monolayer using a gene ontology-defined 250-plex nanostring probe panel

**DOI:** 10.1101/355552

**Authors:** J.M. Robinson, S. Turkington, S.A. Abey, N. Kenea, W.A. Henderson

**Author notes:** Robinson OrcidID: 0000-0002-1456-2851. Henderson OrcidID: 0000-0003-3924-7118, Scopus Author ID: 32367603100.

## Abstract

The Caco-2 cell line has served a historically important role as *in vitro* model for molecular and cellular biology of polarized intestinal epithelia, including for effects of glucocorticoid hormone Dexamethasone. Glucocorticoid hormones modulate the endogenous stress response and are important pharmaceuticals for inflammatory diseases including IBD, yet while they significantly affect immune cells, less is known about their specific effects upon epithelial cells and specific effect on epithelial permeability. Previous research showed that DEX exposure does not immediately produce a quantitative effect, but only after a prolonged treatment >10 days. Culture age itself causes marked effects in these non-renewing cell layers which acts as a confounding variable for observed DEX results. To improve resolution of GC-responsive gene expression in this context, we tested polarized Caco-2 monolayer cultures during at 30-day timecourse, with ~15-days of continuous Dexamethasone exposure. We tested differential gene expression using a 250-plex gene expression panel with the Nanostring nCounter^®^ system, with multiple replicates collected periodically over the timecourse. Gene panel was selectively enriched a-priori for KEGG pathway annotations from tight-junction, actin cytoskeleton regulation, colorectal cancer pathways and others, allowing highly focused, gene-set pathway enrichment analyses. Nanostring nSolver™ data modelling algorithm uses an optimization algorithm and mixture negative binomial model to factor for Time and DEX covariate effects during determination of DE. Analysis identifies strong, culture age-associated “EMT-like” signature with upregulation of actomyosin contraction and integrins, while DEX treatment is associated with a subtler, yet significant counter-signal with suppression of actomyosin genes, and selective DE for different RTKs.

## Introduction

Glucocorticoids modulate stress response in tissues throughout the body including perturbation of gastrointestinal barrier function associated with intestinal inflammation [1]. Immunosuppressive effects of GCs make them important pharmacological agents against chronic disorders including inflammatory bowel disease [2; 3], while long-term negative side effects of GC use include degeneration of the epithelial barrier [4].

Epithelial barrier function is the result of interactions between the tight-junction (TJ) complex, actin cytoskeleton, and Rho GTPase signaling pathways [5;6]. ZO-1 (TJP) (as well as TJP2, TJP3) proteins act as a hub between the transmembrane proteins and filaments of the actin cytoskeleton; TJ is described as the ‘zonular signalsome’ because of crosstalk between TJ, AJ, and RhoGTPase signaling [7–10]. Trans-membrane protein Occludin (OCLN) prevents paracellular translocation of macromolecules, the Claudin family members provide selective permeability to small ionic molecules [11]. (**Fig.1A**)

**Figure 1.**
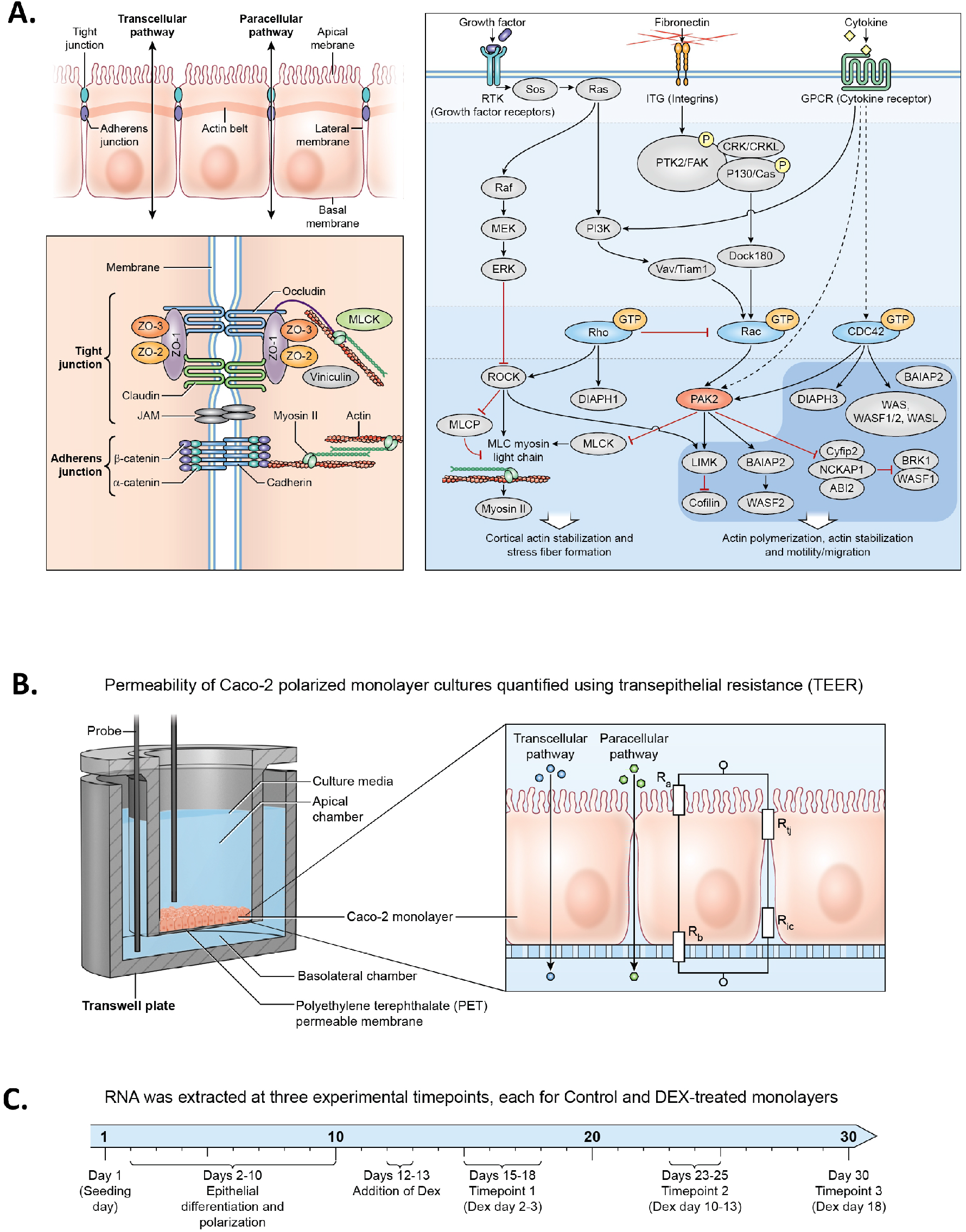
Epithelial identity pathways and experimental design. **1A.** Canonical cellular pathways regulating intestinal epithelial permeability include the tight junction complex, RhoGTPase signal transduction, and and regulation of actomyosin assembly. **1B.** Caco-2 monolayers are cultured on a permeable membrane system, facilitating measurement of alterations in trans-epithelial permeability using TEER. **1C.** Timeline for the two experiments: 1) D1-D30 timecourse with 1 time variable. This timecourse covers the full period of seed population proliferation, epithelial polarization, and effects of 30-day ageing of the cell culture. 2). D15-D30 2 variable, Dexamethasone vs. Control. This timecourse covers effects of DEX over 15 days starting with fully polarized epithelial monolayer.

The Caco-2 cell line has a decades-long history of use as *in vitro* model for pharmacokinetic assays for drug absorption [12]. Recent papers used the Caco-2 cell line for investigations of regulatory mechanisms of protein and mRNA expression of TJ proteins during stimulation with inflammatory cytokines and/or treatment with GCs [13–16]. Two reports specifically investigated GC regulatory effects during long-term timecourse on epithelial polarization experiments (>21 days) [17, 18]. These papers showed GC effects occurring significantly only at later stages of the timecourse and indicate that GC may have important biological effects only after multiple weeks of DEX exposure. Microarray datasets for long-term Caco-2 polarization timecourse have been published as well, which show significant expression of tight junction members (available via the NCBI GEO database) [19, 20].

Questions we sought to answer were: **1)** How are gene expression and pathway-associated gene sets affected during a 35-day timecourse? **2)** Given a 20-day DEX treatment, what is the relative DEX-associated vs. timeline-associated differential gene expression signal? We implemented a medium-throughput gene expression assay using nCounter™ system with Caco-2 monolayers grown on apico-basolateral membrane system and collected trans-epithelial resistance (TEER) as a readout for permeability (**Fig.1B**). We grew cells for 30-days, with DEX treatment between days 15-30 (15 days) (**Fig.1C**).

Our methodology represents some innovation relevant to scale-up of a small-laboratory gene expression workflow. All expression data was obtained from a partially automated, medium-throughput gene expression platform with integrated bioinformatics pathway analysis, the Nanostring nCounter^®^ system and nSolver™ software. Nanostring methodology utilizes fluorescent barcode-labelled hybridization probes, does not use reverse-transcription but provides a single molecule, optical counting method providing key technical advantages over both RNA-seq and qPCR methods [21]. Our custom 250-plex probe panel selected genes from canonical KEGG pathways of interest for intestinal epithelial permeability function – Tight Junction, Regulation of Actin Cytoskeleton, and Colorectal Cancer pathways [22, 23]. Our analysis utilized pathway scoring as a data reduction method, and gene-set enrichment analysis for testing statistical support for prospectively enriched pathways. Pathway scoring was performed using Reactome pathway database annotations, a new database with significant functional pathway data [24].

Using this workflow, we infer molecular-cellular functional effects associated with culture age and DEX treatment using a medium-sized gene expression panel, with good statistical support for pathway enrichment. Further development of this workflow seeks to implement additional automated functional readouts for the workflow, such as protein expression and co-localization, automated TEER data collection, cell cycle/cellular proliferation assays, and utilization additional statistical methods for multivariate regression and machine learning.

## Materials and Methods

### Caco-2 cell culture

Caco-2 cell line was purchased from American Type Culture Collection (ATCC# HTB-37), and cultured consistent with best practices specified in the ATCC manual (https://www.atcc.org/Products/All/HTB-37.aspx?geo_country=us), and reported in literature. and basic research in the molecular and cellular mechanisms of epithelial barrier function [12, 25]. Cryovials were thawed, resuspended, and pelleted to remove DMSO and suspended in 12mL in T75 flasks. Cells were grown to 60-80% confluency and split by aspirating media, rinsing with HBSS, and harvested with .25% Trypsin-EDTA and split 1-2 or 1-3. Viable cell counts were obtained using Trypan Blue (BioWhittaker^®^ catalog# 17-942E) and Nexcelom Bioscience Cellomenter™ Auto T4 cell counter. Cells were initially thawed from cryopreserved vials with 5 or 6 previous passages, seeded after expanding for 3-4 additional passages.

Cells were seeded onto 24-well insert plates with PET) permeable membranes with 1.0 uM pore size (Millipore product # PSRP010). Seeding volume was 300uL cells at density of .4×10^6^ cells/mL (or 120,000 cells/well) in the apical chamber. Cultures were grown at 37°C, 5% CO2 on filter-sterilized Eagle’s Minimum Essential Medium (EMEM) with 20% Fetal Bovine Serum, with Penicillin-Streptomycin (Quality Biological™ cat. # 120-095-671). Media was changed every other day during the experiments.

### Trans-Epithelial Electrical Resistance (TEER)

TEER readings were taken at regular intervals during the experimental timecourse with the Millicell ERS-2 Epithelial Volt-Ohm Meter (cat. # MERS00002). Cell cultures were equilibrated at room-temperature (22°C) before readings were taken, to control for temperature effects on TEER readings [26].

### Timecourse and DEX treatment

Dexamethasone treatment began ~15 days post-seeding, at which point epithelial monolayers become polarized, after 2-3 days of stable cell density (**Fig.2B**). DEX was applied to experimental wells at 10uM, with one set of replicates treated at 100uM. Sigma reagent D2915 “Dexamethasone-Water Soluble” was used. Water-soluble DEX is compounded with methyl-β-cyclodextrin, product documentation states the dry reagent is 65mg/g DEX by weight. DEX reagent was labelled 392.5g/mol. Stock solution was made by dissolving DEX at 10 grams/L. 10uM Dex in media was obtained by using: 65mg/g x 10g/L = .65g/L x 1mol/392.5g = .00166mol/L = 1.66mM = 1660uM. For 10uM, added .06mL Dexamethasone-Water Soluble stock to 10mL media. Control cell cultures were grown in standard media without DEX or carrier. As DEX is commonly orally administered in conjugated form, the experimental treatment can be interpreted as a model for cellular action during pharmacokinetic exposure of conjugated Dex *per se*, rather than generalized to glucocorticoid activity in general. Total RNA was extracted from fresh monolayers at multiple timepoints during the full 30-day timecourse, and during the 15-day DEX treatment; the two timecourses were analyzed as separate experiments (**Fig.2A, Fig.3C, respectively**).

**Figure 2.**
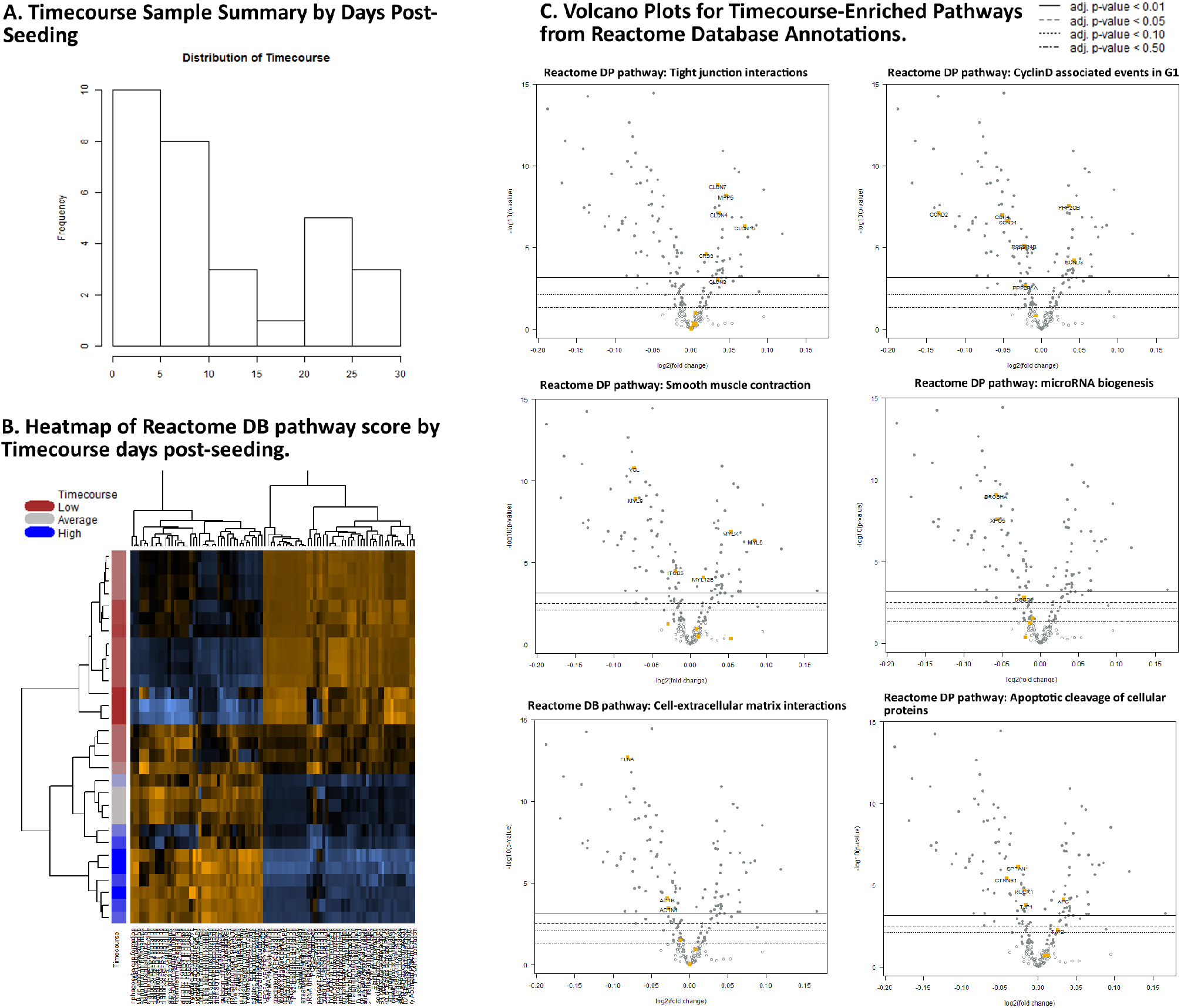
D1-D35 Caco-2 Timecourse, ‘Cellular Ageing’ Experiment. **2A.** Untreated sample replicates (technical and biological) per timepoint, used in this analysis. Histogram shows sample number collected for each binned timepoint. **2B.** Heatmap of Pathway Scores, using ReactomeDB pathway annotations for genes. Clear separation of pathway activity is observed associated with early- and later-stage cultures. **2C.** Volcano plots showing differentially expressed genes by their associated ReactomeDB annotations, for pathways with statistically significant differentials under Gene Set Analysis. For example, many Claudins have significantly increased DE over time, each Claudin has a ReactomeDB “Tight Junction Interactions” annotation.

**Figure 3.**
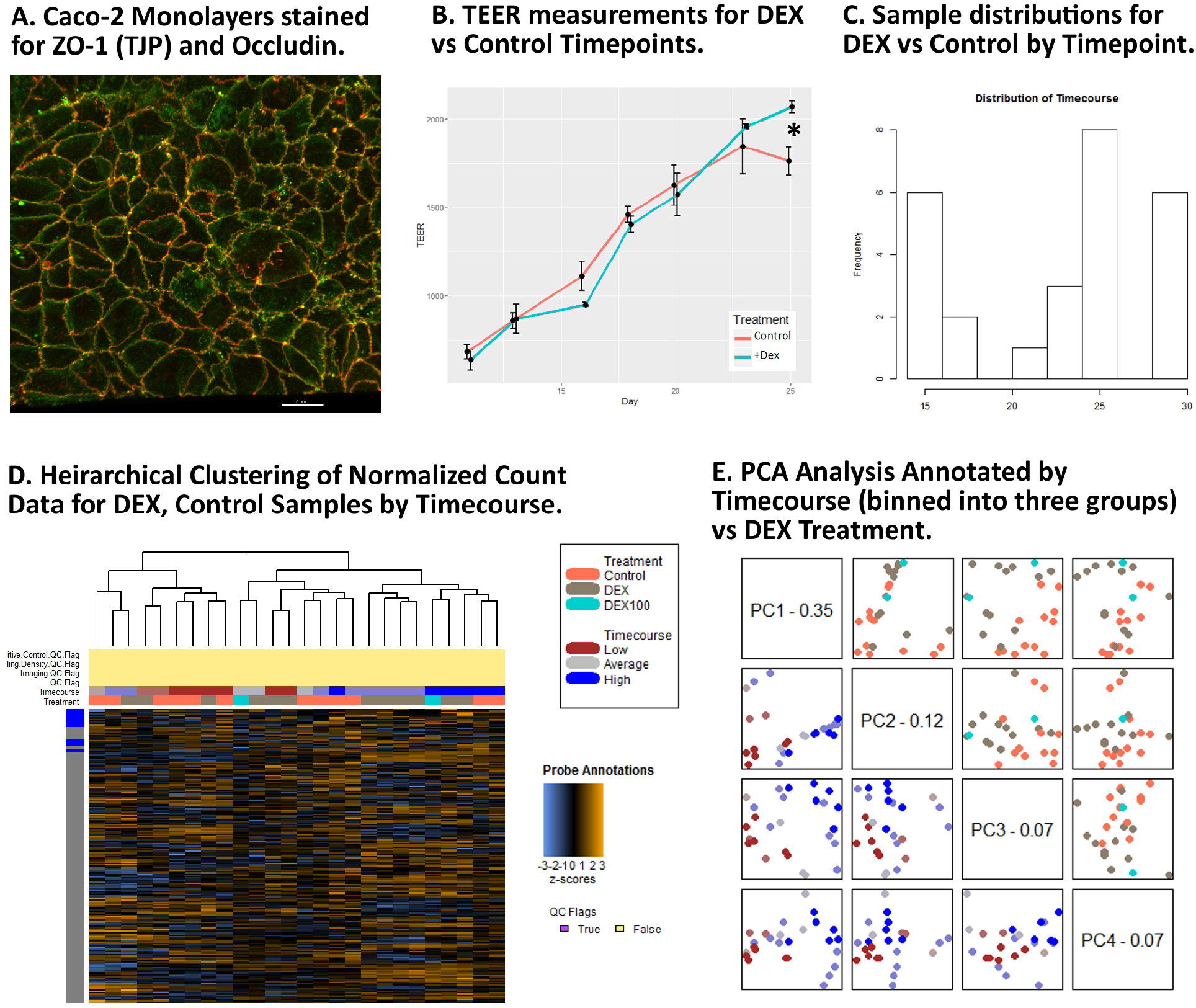
Dexamethasone vs. Controls Timecourse. **3A.** Confocal image of Caco-2 polarized monolayer (at ~10 days, immunofluorescent staining for Occludin and ZO-1). **3B.** Trans-epithelial electrical resistance (ohms/cm^2^) average readings for DEX and Control samples during the timecourse. As oberseved in previous research, DEX does not affect TEER until ~15 days after treatment starts. **3C.** Distribution of sample numbers used in this analysis by days post-seeding. **3D.** Unsupervised clustering of normalized gene expression for the full gene panel. **3E.** PCA results showing the distribution of variable sets by principle component. PC1 is reasonable predictor of timecourse progression, PC3 is a marginally effective predictor for DEX treatment though neither perfectly separate the respective variables.

### Experimental Replicates

We collected RNA samples for 2-4 plate replicates each from 2-3 biological replicates for selected timepoints both the 30-day timecourse and the Day15-30 DEX treatment timecourse. Replicates were collected at various days, with days after seeding treated as a continuous time variable during differential expression testing.

### RNA Extraction

Total RNA including small RNA from Caco-2 monolayers was extracted from individual permeable membrane inserts at the three experimental timepoints using the Qiagen miRNeasy^®^ Micro kit (cat. #217084) with RNeasy^®^ MinElute^®^ spin columns (mat. #1026497). Extracted total RNA was quantified with Thermo Scientific™ NanoDrop™ spectrophotometer. RNA was stored at −80°C until expression testing.

### Nanostring expression profiling and probe-panel selection

Our probe panel of 250 gene targets was designed to leverage the advantages of the Nanostring nCounter^®^ and nSolver™ software. Complete probe sequences and isoform coverage for the panel are provided (**Sup. Methods 1.1, Probe Panel Specifications**); also see Robinson and Henderson, 2018[23] for further description of panel selection. Briefly, our gene selection strategy used KEGG pathway database [22] to generate a list of genes affecting intestinal-epithelial barrier function and epithelial homeostasis. Pathways selected from included tight-junction (map04530), regulation of actin cytoskeleton (map04810), colorectal cancer (map05210), most with significant overlapping membership in adherens junction (map04520), focal adhesion (map04510), Wnt (map04310), and other pathways (Supplementary Methods available at publication). Genes from initial list were excluded if not expressed in lower GI tissues by referencing against the Human Protein Expression Atlas (which includes tissue-specific RNA expression) [27]. Additional genes of interest were included on an ad hoc basis:

1. KEGG microRNA gene targets associated with colorectal cancer (map05206). These include Programmed cell death 4 (PDCD4), DNA methyltransferase 3A (DNMT3A), and others.
2. Core microRNA biogenesis pathway genes Drosha (RNASEN), Pasha (DGCR8), Dicer, Argonaute2, Exportin5, and the RISC loading complex RNA binding subunit (TRBP).
3. Putative reported biomarkers for GI dysfunction [28] including chemokine receptor CXCR4, Lysozyme (LYZ) and others.
4. Histological enterocyte differentiation markers Carbonic Anhydrase I and II (CAI, CA2) [29], and cancer stem marker Aldehyde dehydrogenase 18, isoform A1 [30].
5. Additional genes with high specific expression in gastrointestinal tissue according to Human Protein Expression Atlas, including additional Claudins, heat shock proteins, and others.

### Differential expression

Analysis of read-count data includes imaging QC, read-count QC, differential expression, and pathway analysis were performed using the Nanostring nCounter nSolver™ 4.0 (Nanostring MAN-C0019-08) with Nanostring Advanced Analysis Module 2.0 plugin (Nanostring MAN-10030-03), following the Nanostring Gene Expression Data Analysis Guidelines (Nanostring MAN-C0011-04). The computational methods are extensively described in these user manual documents.

Advanced Analysis Module 2.0 software uses open-source R for QC, normalization, differential expression analysis, pathway scoring, and gene set enrichment analysis. R-code is provided together with complete results of the bioinformatic workflow in an html-format. All QC, normalization, differential expression, and pathway analysis data and graphical outputs are available upon request prior to peer-reviewed acceptance.

Normalization reference genes were obtained from geNorm algorithm [31] implemented in the R-language Bioconductor package NormqPCR [32]. geNorm selects reference probes based on global stability (M_i_) of pairwise expression ratio between samples, iteratively identifying genes with the lease expression variance, we chose 10 genes with minimum M_i_ values as normalization reference genes. Normalized probe count data for each experiment is provided (**Sup. Results 1.1, Normalized Counts**)

The data model preferentially applies the optimal statistical method per gene given the variable distribution 1) Mixture negative binomial model, 2) Simplified negative binomial model, 3) Log-linear model, in that order, for determination of DE. FDR p-value adjustment was performed with Benjamini-Yekutieli method. Under the data model, time (day post seeding) was used as a continuous variable for to testing DE during the D1-D30 timecourse. As predictor variables for the D15-D30 DEX treatment experiment, time (day post seeding) as a continuous variable, and Control/DEX+/DEX100 were used as categorical variables with Control as the reference category. All analyses were performed with batch and cartridge IDs factored as confounding variables in the data model and exerted minimal effect on the DE determinations. (**Sup. Results 1.2, 1.3, 1.4, DE results for each variable comparison**)

### Pathway scoring and gene-set analyses

Pathway scoring and gene-set analysis were performed to gain statistical support for pathway functional enrichments using the Nanostring nSolver Advanced Analysis Module 2.0 application. Analyses are based on open-source R packages, with open-source code provided as part of the supplementary materials. Pathway scores are derived by calculating the first principle component of pathway genes’ normalized expression. (**Sup. Results 2.1, 2.2, Pathway scoring results for experimental comparisons**)

Gene set analysis is a quantitative summary of differential expression for gene sets (gene annotations equivalent to those for pathway scoring). GSA summarizes the differential expression for genes from each annotation, then calculates a global differential significance score for each gene set, allowing for some quantitative inference of putative functional effects. (**Sup. Results 3, GSA global statistics**)

### Database accession

All raw and normalized Nanostring count data are pending submission and release to NCBI Gene Expression Omnibus (GEO) database upon publication of the article. Sample data and replicate information have been accessioned in the NCBI GenBank BioProject and BioSample databases, available pending publication (BioProject ID: PRJNA525237).

## Results

Our primary objective in these experiments was to identify signal from differentially expressed DEX-associated genes against a high background of culture age-associated effects, secondarily to describe differential expression for the entire, D1-D30 timecourse. In this version (Robinson et al. v2), we identified that differentially expressed genes were very similar to those identified in Robinson et al. v1. Changes in the experiment included using time variable as a continuous, rather than categorical variable as was done previously. We also performed additional biological replicates across all timepoints, added additional biological replicates, and implemented a more precise statistical testing and optimization algorithms. Finally, we compared our results with the results of microarray analysis from a similar Caco-2 timeline [19, 20], and found that expression patterns described there were concordant with differentially expressed genes from our probe panel subset.

### Experiment 1: Differential expression associated with 30-day timecourse

During the D1-D30 timecourse, many differentially expressed genes were identified, using day-post-seeding number as a continuous variable. Pathway scoring results in a strong pattern of differential expression with inflection point roughly between low-medium ranges, around D10 (**Fig.2B**). This corresponds roughly to culture time previously published reports and experiments have used for complete Caco-2 polarization. ReactomeDB annotated gene-set enrichment analysis provides some indication of directionality over time for several key epithelial pathways base on increased vs. decreased expression of pathway consituents. Over time for example, many Claudins show increased expression for the ‘Tight junction interactions’ and ‘Apoptotic cleavage of cellular proteins’ pathways, while ‘Cyclin-D associated events in G1’, ‘miRNA biogenesis’ pathway genes are mostly decreased in expression, while others show variable increase and decrease of gene expression in several multiple pathways with enrichment of differentially expressed constituent gene members (**Fig.2C, Supplementary Results available at publication**).

### Experiment 2: Differential expression associated with Dexamethasone

Our second experiment was consisted of the D15-D30 DEX treatment, using days-post-seeding again as a co-variate, with just this later set of timepoints, to capture events occurring after polarization. TEER increased steadily during the timecourse, indicating steadily decreasing permeability as the Caco-2 monolayer forming an increasingly less permeability in both DEX treated and control cell cultures. Between culture days 23-25 however (~D13-15 of DEX continuous exposure), DEX-treated cultures showed significantly higher TEER relative to control cultures (**Fig.3B**), a pattern is consistent with previously observed results of Fischer et al. (2014) and Zheng et al. (2017) [17, 18].

Consistent with this, differential pathway scores for DEX-treated samples showed stronger clustering at later timepoints, where at earlier timepoints, DEX-treated samples cluster within Control groups. (**Fig.4B**) Pathway scores shown as a heatmap of ratios between the experimental comparisons show that while there is some overlap between effects of time and DEX variables, each of the DEX vs. Control and Timecourse variables have pathway enrichment specific to that variable (**Fig.4C**).

**Figure 4.**
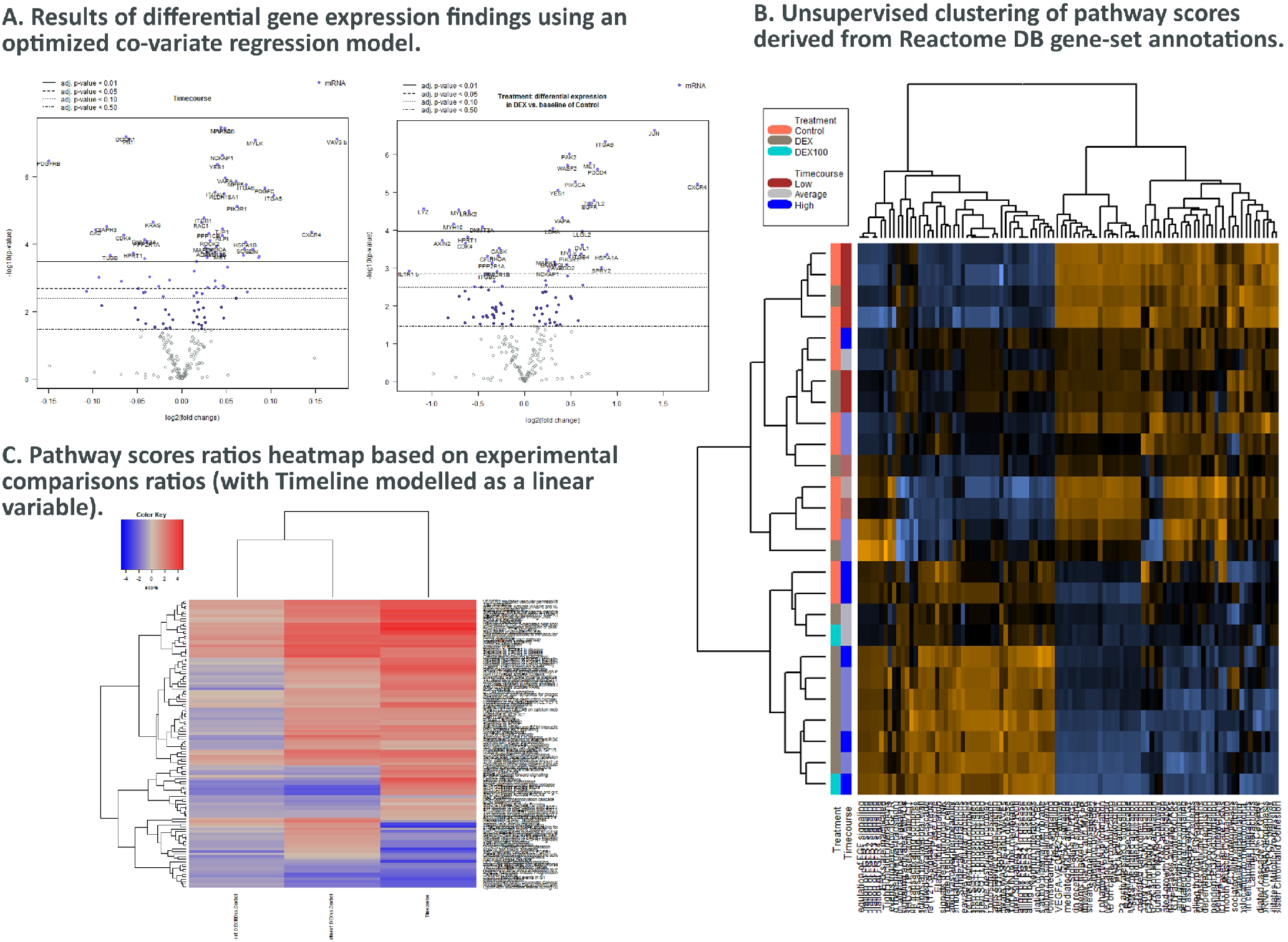
Dexamethasone vs. Controls Timecourse. **4A.** Volcano plots for each respective covariate. **4B.** Clustering of raw pathway scores. **4C.** Clustering of differential pathway scores for each experimental comparison.

Two of the key pathways exhibit opposite behavior respective of the variables Time and DEX. These include the ReactomeDB ‘Rho GTPases Activate ROCKs’ pathway, which has nearly opposite expression patterns in the respective variables (**Fig.5A,5C**). In the ‘Apoptotic cleavage of cellular proteins’ pathway, which includes tight-junction Occludin and ZO-1, there is insignificant differential expression associated with DEX, but increased pathway score associated with the Time variable (**Fig.5B,5D**). Enterocyte differentiation marker Carbonic anhydrase 2 (CA2) while the colorectal cancer stem marker Aldehyde dehydrogenase 18 A1 (ALDH18A1), pattern is consistent with Caco-2 monolayers developing diminished epithelial differentiation and with increasing tendency towards ‘stemness’ over time.

**Figure 5.**
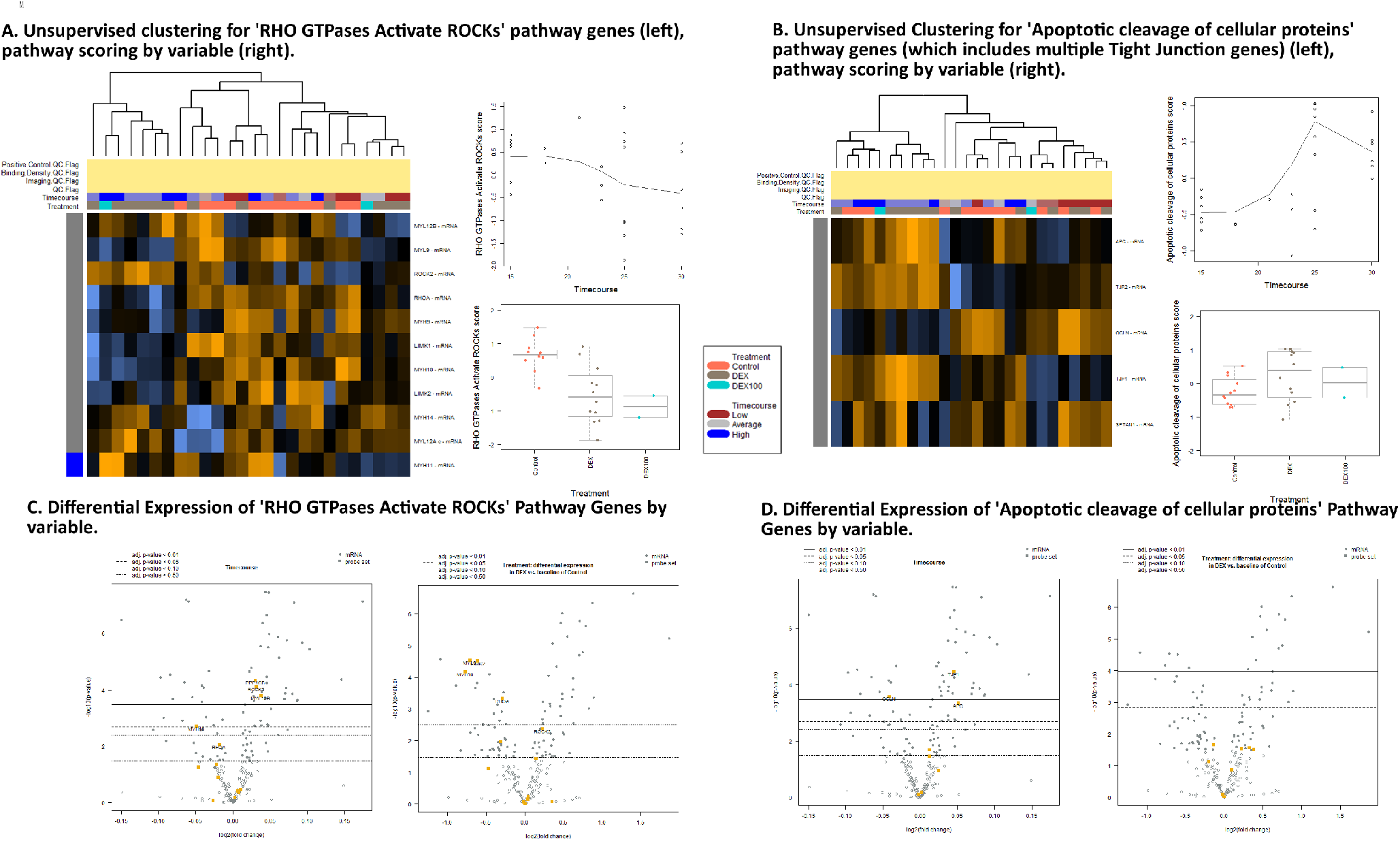
Pathway score and GSA results for Dexamethasone vs. Controls Timecourse. **5A,C.** Expression of genes associated with RHO GTPases Activate ROCKs ReactomeDB pathway show activity associated with DEX treatment. **5B,D.** Expression of genes associated with Apoptotic cleavage of cellular proteins pathway shows activity associated with timecourse, but not with DEX treatments.

Overall, there is less differential expression associated with DEX than the expression occurring over time during the process of cellular ageing in terms of both p-values and fold changes, however the advanced statistical modelling of co-variates provides well-supported DEX-associated differential expression results as well. The KEGG Pathview) [33] for the ‘Regulation of Actin Cytoskeleton’ pathway, shows that generally, DEX treatment is associated broadly with increased expression of cell surface cytokine receptor CXCR4, integrins, epithelial and hepatocyte growth factor receptor tyrosine kinases. Diminished expression of acto-myosin assembly and regulatory components and PDGFR- and FGFR-TKRs (**Fig.6**) are observed.

**Figure 6.**
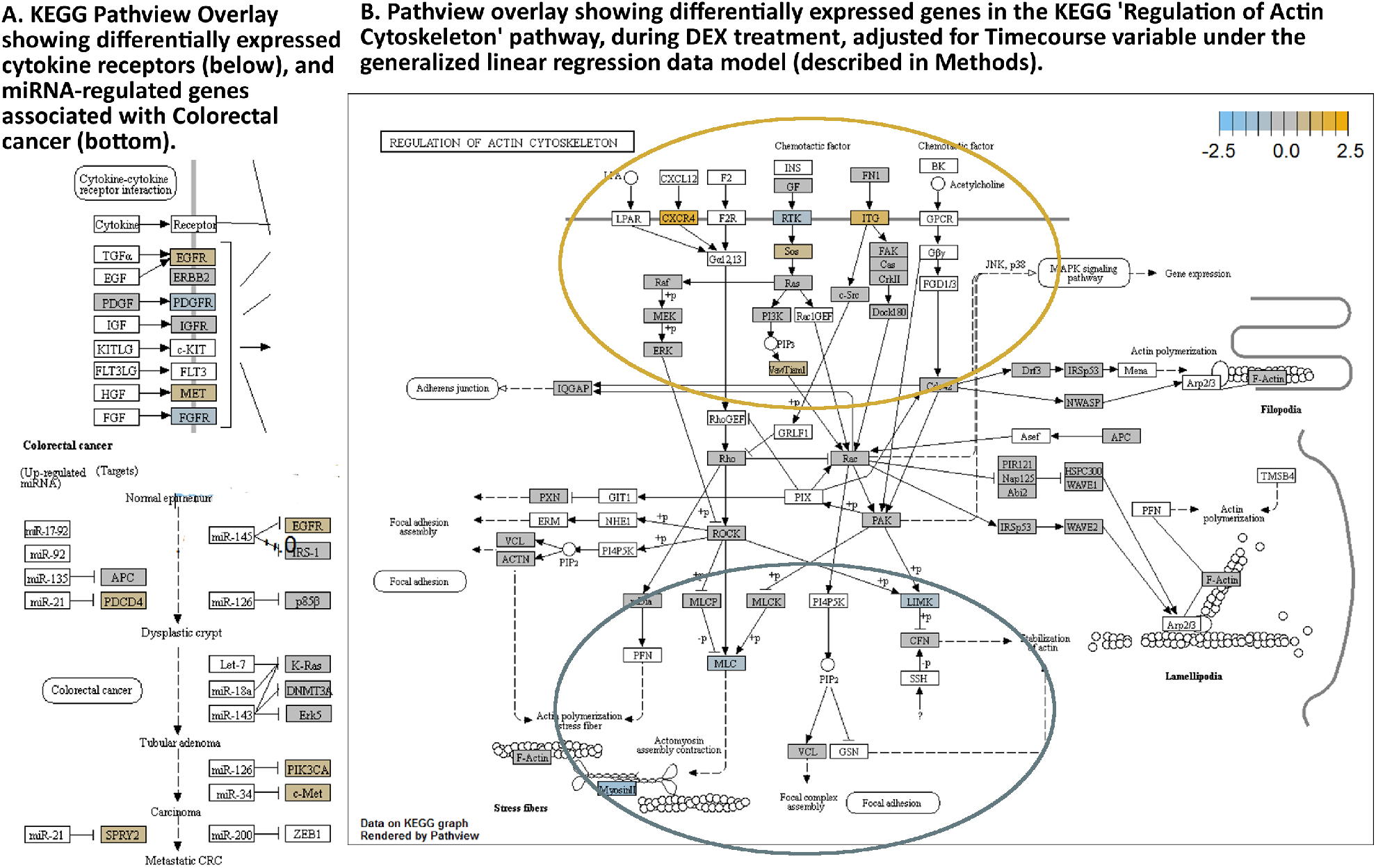
KEGG Pathview readouts of DEX vs Control-associated differential gene expression. **6A.** Differentially expressed cytokine receptors and known miRNA targets in colorectal cancer. **6B.** Pathway map for KEGG “Regulation of Actin Cytoskeleton” shows effects of DEX treatment on cellular pathway components. (white backgrounds not assayed)

## Discussion

### Increased permeability in D15 DEX vs. Controls

Observation of increasing TEER values over time indicates permeability decreasing over time in maturing Caco-2 cultures. The trend of increasing TEER continues from initial seeding up to approximately culture day 25 (~Dex-day 15). At this point, TEER in DEX-treated and control cultures become significantly divergent: DEX-treated cultures continue to exhibit increasing TEER/lowered permeability, while in Controls, TEER levels off (**Fig. 3B**). These results are consistent with previous reporting of similar result [17, 18]. Observable effects on permeability from DEX treatment do not appear to become significant until at least 20-25 days post-seeding, long-term effects of GC exposure should therefore always be considered when utilizing long-term cultures of this *in vitro* model. Occludin showed significantly lowered expression over time, while ZO-1 (TJP) increases, but is not significantly associated with DEX treatment (**Fig.5B,D**).

### Dexamethasone-associated differential gene expression

DEX-treated cultures exhibit a specific decrease in actomyosin stress-fiber assembly/contraction pathway genes including Myosin 10 (MYH10), its regulator Myosin light-chain regulatory peptide 9 (MYL9), the negative regulator of actin stabilization LIM-domain kinase 2 (LIMK2), and calcium/calmodulin-dependent serine/threonine kinase (CASK, associated with membrane trafficking and centrosome formation) (**Fig.5**). The lowered expression may result in inhibition of stress-fiber formation and alteration of the cortical actin cytoskeleton by DEX, which could provide some suppressive effects against a full EMT progression [34].

### Putative biomarkers for GI pain and inflammation

Lysozyme (LYZ) is a biomarker for GI inflammation [35], shows showing significantly lower expression in our DEX-treated cultures (**Fig.4A**). LYZ is associated with mucous/sodium flux and microbial dysbiosis associated with stress response. CXCR4 is a leukocyte trans-endothelial migration associated cell surface receptor, which has been associated as a putative IBS-associated biomarker [28].

### DEX-associated c-Jun expression

DEX-associated increase in expression of the pro-survival transcription factor c-JUN (**Fig.4A**) is interesting in light of previous reports that GCR/AP-1 promotor binding suppresses JUN transcription [36]; this may be a phenomenon resulting from adenocarcinoma-specific alterations present in the cell line. However, DEX-associated, *ras*-dependent stimulation of c-Jun expression in the rat intestinal epithelial cell line IEC-6 was reported by Boudreau et al. (1999) providing some indication that DEX-associated c-Jun activation may potentially be an intestinal-epithelial specific response [37].

### Dexamethasone carrier molecules may have significant effects

Dexamethasone is not water soluble, so is often compounded with a carrier such as a Cyclodextrin molecule which increases the solubility (and bio-availability), of the Dexamethasone [38, 39]. Our data should be interpreted as the result of Dexamethasone-Cyclodextrin complex, results are robust to multiple QC and normalization parameters, and are concordant with metadata comparisons as previously described. Carrier effects are an important consideration in pharmacological design and delivery of water-insoluble molecules such as steroid molecules, and understanding the results of Cyclodextrin:Dexamethasone complexes on tissue models is significant for the understanding pharmacological action [40]. Beig et al. (2013) and Fine-Shamier et al. (2017) report cyclodextrin-based formulations to result in decreased permeability, or pericellular transport, of DEX however DEX should be bioavailable for the epithelial monolayer [41, 42].

### Future directions

This workflow proved useful for understanding co-transcriptional responses in an efficient, medium-throughput format. Transcriptomics projects often face large, noisy datasets for which relevant functional signal may be rather subtle, a long-time strategy for addressing this has been to focus analyses on specific gene subsets rather the global transcriptome. A wide variety of novel approaches has been developed for post-hoc dimensionality of such data, for example[43], though less explored are the best practices for a-priori, pathway-based panel selection. We Focus on just 250 genes from relevant pathways eliminated the need for more intensive bioinformatics necessary for RNA-seq projects, allowed us flexibility with ad-hoc gene additions, and a high degree of responsiveness in selecting technical and biological replicates. Follow-up projects which we have utilized this system (in-prep) include comparison of multiple GI epithelial cell types, cross-species comparisons, and transfection of microRNA mimics.

## Supporting information

Supplementary Results 1

Supplementary Results 2

Supplementary Results 3

## Abbreviations

(GCs): Glucocorticoids
(GR, geneID hsNR3C1): Glucocorticoid Receptor
(GREs): Glucocorticoid Response Elements
(OCLN): Occludin
(TJP, ZO-1): Zona Occludans-1
(DEX): Dexamethasone
(CRC): Colorectal Cancer
(TJ): Tight Junction
(IECs): Intestinal Epithelial Cells
(AJ): Adherens Junction
(TEER): Trans-epithelial (electrical) resistance
(KEGG): Kyoto Encyclopedia of Genes and Genomes
(FDR): False Discovery Rate

## Author Contributions

**Robinson** designed the experiments, conceived and designed the custom Nanostring codeset, performed cell cultures, collected TEER data, performed RNA extractions, performed Nanostring protocols and operated the nCounter instrument for each replicate, on-site at the Henderson Lab in the NIH Clinical Center. Robinson performed statistical and bioinformatic analyses using the R-based nSolver™ 4.0 and Advanced Analysis 2.0 software packages, with manual checking of results, and TEER data analysis using R/R-Studio. Robinson wrote and revised the version 2 text and produced figures (see acknowledgements for additional Figure 1 and 2 contributions).

**Turkington** performed cell culture tasks, RNA extraction, and operation of Nanostring nCounter^®^.

**Abey** initially developed cell culture methods, contributed to Nanostring codeset design and content, and provided comments for the text.

**Kenea** performed additional cell culture tasks.

**Henderson** designed the experiments, contributed to the Nanostring codeset design and content, organized and supervised research and administration, edited and revised the manuscript.

## Acknowledgements

Robinson, Turkington, Kenea were funded through NIH IRTA fellowships. Research was funded to Wendy A. Henderson through 1Z1ANR000018-01-8. Authors acknowledge Ethan Tyler of the NIH Medical Arts Division for his graphic design work on Fig.1, and Dr. Greg Gonye for Nanostring nCounter training.

## Disclosures

All authors declare no conflicts of interest.

